# Cardiovascular responses to natural and auditory evoked slow waves predict post-sleep cardiac function

**DOI:** 10.1101/2024.05.03.592377

**Authors:** Giulia Alessandrelli, Stephanie Huwiler, Giulio Bernardi, Manuel Carro-Domínguez, Fabia Stich, Rossella Sala, Florent Aziri, Anna Trippel, Susanne Markendorf, David Niederseer, Philipp Bohm, Pietro Cerveri, Francesca Siclari, Reto Huber, Nicole Wenderoth, Christian Schmied, Caroline Lustenberger

**Author notes:** Corresponding Author: Caroline Lustenberger, Neural Control of Movement Lab, Department of Health Sciencesand Technology, ETH Zurich, 8092 Zurich, Switzerland. these authors contributed equally to this work (Giulia Alessandrelli, Stephanie Huwiler).

## Abstract

The interplay between slow-wave sleep and cardiovascular health is increasingly recognized. Our prior research showed that auditory-enhanced slow waves can boost cardiac function, yet the mechanisms behind this remain unclear. Advancing these findings, our current analysis dissected the effects of two slow wave types on cardiovascular function, using data from 18 middle-aged men across three nights. We found that the strength of heart rate and blood pressure responses concurrent with slow waves predicts cardiac function post-sleep. Notably, we identified that highly synchronized type I slow waves, as opposed to lower-amplitude type II slow waves, primarily co-occur with these cardiovascular pulsations. While auditory stimulation enhances both types of slow waves, they exhibit distinct temporal dynamics, pointing to different underlying biological mechanisms. This study crucially addresses how distinct slow wave types can affect cardiovascular function, implying that targeted slow wave stimulation could be a strategic approach to improve heart health.

## Introduction

With humans spending approximately one-third of their lives sleeping, sleep’s implications influence the entire spectrum of health and disease. The scientific community has increasingly recognized the bidirectional relationship between sleep and cardiovascular function, with changes in sleep macro-architecture being both a consequence and a contributor to cardiovascular diseases ^1–4^.

The deepest stages of non-rapid eye movement (NREM) sleep seem to be crucial for cardiovascular recovery and health. Specifically slow waves, the hallmark oscillations of deep NREM sleep, may play a critical role in promoting cardiovascular function during sleep. Auditory stimulation during NREM sleep is a feasible approach to effectively modulate sleep slow waves ^5–9^. However, the influence of auditory stimulation beyond modulating cortical activity, such as its impact on the autonomic nervous system, which could influence cardiovascular function during sleep, remains a topic of ongoing investigation ^9–12^. Our group’s recent findings demonstrated that auditory stimulation enhances slow waves and induces a cardiovascular response, including a transient increase in blood pressure and a biphasic heart rate change, while also increasing post-sleep left ventricular function compared to a control condition ^10^. However, which underlying mechanisms are linked to such cardiovascular benefits remains unclear.

There is growing evidence that two types of slow waves exist, which can be differentiated based on wave morphology, timing, and synchronization processes ^13,14^. Type I slow waves, which exhibit a broad cortical synchronization, are predominantly observed during the early stages of sleep onset and likely result from subcortico-cortical synchronization processes of neurons ^13,14^. In contrast, type II slow waves, resulting from cortico-cortical synchronization, become more prominent during the later stages of sleep onset ^13,14^. K-complexes, which represent the largest deflection observed in EEG readings during NREM sleep and can be triggered by external stimuli or occur spontaneously during sleep, are likely characterized by their substantial neuronal synchronization, potentially via subcortico-cortical synchronization processes, and could thus be considered a specialized subtype of type I slow waves. They have been proposed to play an essential role in promoting cardiovascular homeostasis ^15^ and were linked to a dynamic blood pressure and heart rate response, potentially through a short burst of sympathetic activity concurrent with their occurrence ^15–20^. Given the potential different mechanisms underlying the occurrence of type I and type II slow waves, elucidating whether they display distinct influences on the cardiovascular system is crucial in understanding their implications in promoting cardiovascular health.

Here, we aimed to determine whether these two subtypes of slow waves are associated with distinct changes in cardiovascular activity during sleep. We provide insights into the relationship between the cardiovascular activation occurring during auditory slow wave enhancement and post-sleep cardiac function. Moreover, we show that previously proposed distinctive types of slow waves based on cortical features are also associated with different cardiovascular dynamics, with type I slow waves evoking stronger cardiovascular responses compared to type II slow waves. Furthermore, we demonstrate that auditory stimulation differently evokes type I or type II slow waves during times of stimulation, leading to important insights on how to apply auditory stimulation to result in cardiovascular benefits the next morning.

## Results

To identify specific mechanisms of how sleep can promote post-sleep cardiac function we examined the differential impacts of type I and type II slow waves on sleep and post-sleep cardiovascular function. We report the results of 18 healthy middle-aged male participants all undergoing three experimental nights in a sleep laboratory according to a randomized-controlled, cross-over study design as illustrated in Figure 1 ^10^. However, one participant was completely excluded because of technical failure in sleep EEG data acquisition. During these sessions, participants were subjected to one of three conditions: SHAM stimulation (no auditory slow wave stimulation), auditory stimulation at 45 dB (*ISI1*_High_), or auditory stimulation at 42.5 dB (*ISI1*_Low_) to enhance slow waves while EEG, ECG, and blood pressure were continuously recorded. Acknowledging that type I slow waves, including K-complexes, can be characterized as highly synchronized slow waves with subcortical origin, whereas type II slow waves are smaller slow waves of cortical origin, we used an established algorithm to automatically separate these slow wave types across the whole sleeping period ^13,14^. We first investigated whether auditory stimulation induces cardiovascular dynamics during sleep compared to a SHAM control condition. Afterward, we established specific characteristics of the heart rate and blood pressure responses, here referred to as cardiovascular response, that predict changes in post-sleep cardiac function assessed through transthoracic echocardiography (TTE). Furthermore, we explored how the cardiovascular response differs between type I and type II slow waves during sleep and how auditory stimulation evokes both types of slow waves with different probabilities. Altogether, we provide insights into the cardiovascular correlates of the two distinctive types of slow waves and how their associated cardiovascular dynamics during sleep are related to cardiac function after sleep.

**Fig. 1:**
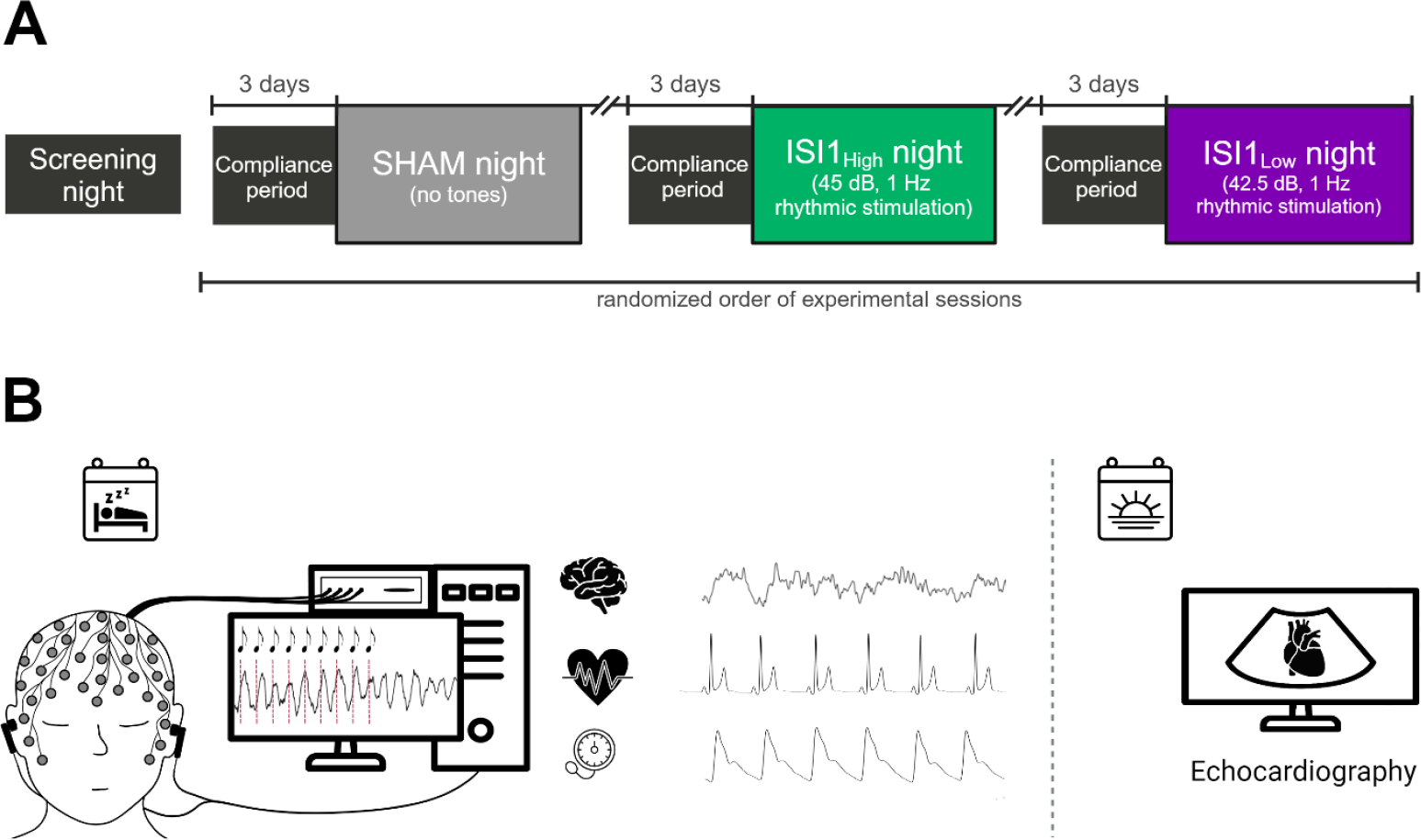
Study protocol of clinical trial. A) First, there was a screening period where the eligibility of participants was checked by an initial phone screening followed by a screening night in the sleep laboratory. If all inclusion/exclusion criteria were fulfilled, participants were invited to a cardiovascular screening to verify the absence of cardiovascular diseases. Afterwards, the experimental phase including three experimental nights took place. Three days before each experimental night, a compliance period started where participants had to adhere to a regular sleep rhythm. The intervention condition was administered in a randomized order. B): Overview of here reported measurements during the experimental night. Participants got a 7.5h sleep opportunity where polysomnography, electrocardiography, and continuous blood pressure were recorded. During sleep, one of the three stimulation conditions according to randomization protocol was applied through an electroencephalography (EEG)-feedback system. In the morning, at the end of each experimental night, an echocardiography recording was obtained by an expert cardiologist.

### Auditory slow wave stimulation induces dynamic heart rate and blood pressure response

To understand the link between auditory enhanced slow waves and cardiovascular dynamics during sleep, we here extracted heart rate and blood pressure from the 2 seconds before to 18 seconds after detected slow wave onsets. Afterwards, we normalized each signal against its average value in the 2-second baseline period. We then calculated the average of these normalized responses for each experimental condition (*ISI*1_High_, *ISI*1_Low_, and SHAM) within individual subjects and across all participants. As seen in Figure 2, there is a significant difference (p<0.05) across experimental conditions, indicating altered cardiovascular responses in both auditory stimulation conditions relative to the SHAM control. Auditory stimulation induced a stronger heart rate response, particularly after 6 seconds of detected slow wave onset, where the heart rate was lowered more strongly. Furthermore, blood pressure showed a temporal increase in stimulation conditions compared to SHAM control.

**Fig. 2:**
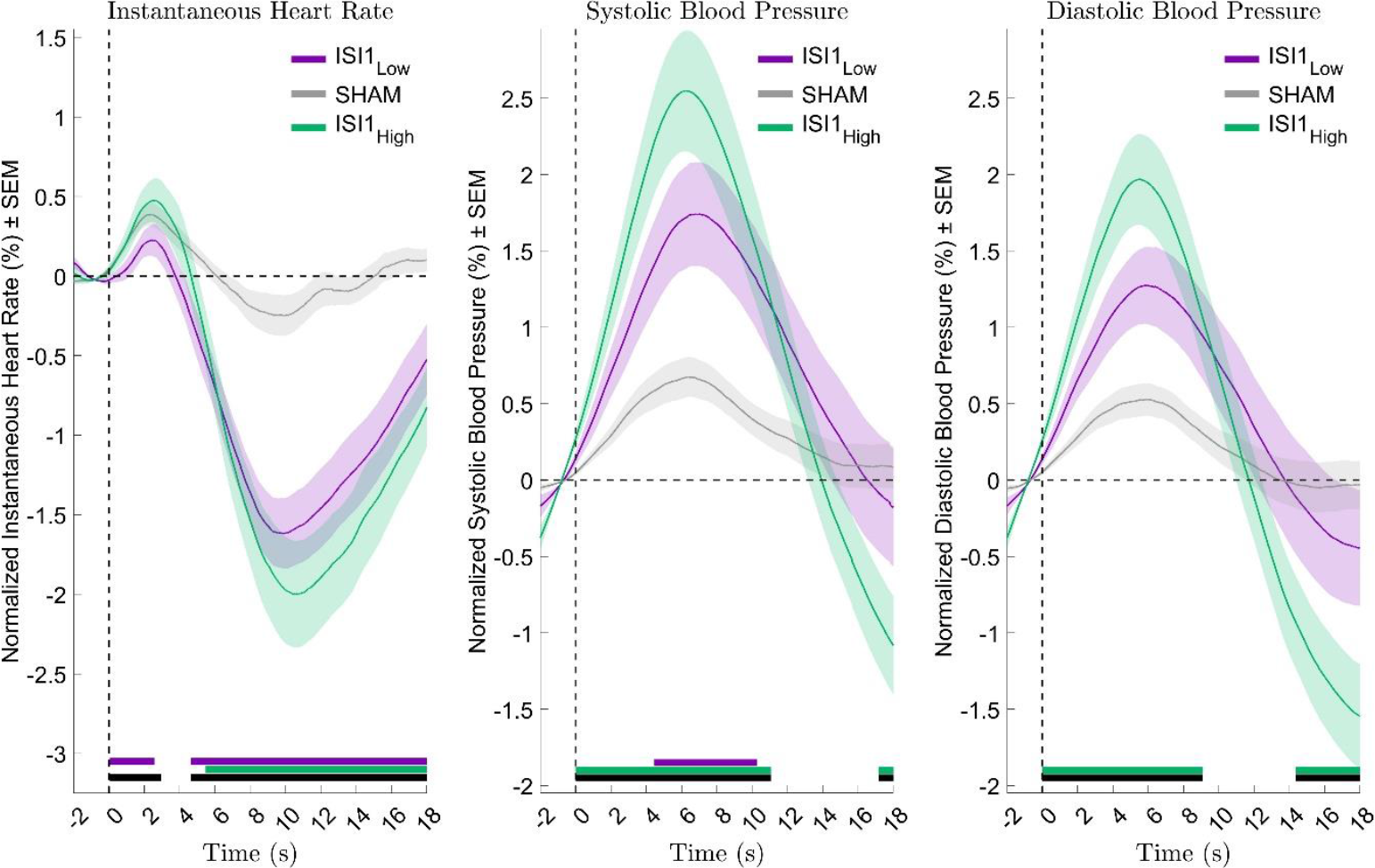
Cardiovascular responses of detected slow waves occurring during stimulation windows. Auditory stimulation significantly affects diastolic and systolic blood pressure and instantaneous heart rate within the three conditions (*ISI1*_High_, *ISI1*_Low_, and SHAM (no tones applied), as indicated through the black bars. The cardiovascular data are normalized against the 2-seconds before the slow wave onset, including 15 for heart rate and 17 subjects for diastolic and systolic blood pressure over three experimental night conditions. Black lines indicate statistically significant (p<0.05) differences between conditions. Purple and green lines indicate significant post-hoc comparisons between auditory stimulation condition and SHAM control with *ISI1*_Low_ and *ISI1*_High_, respectively.

### Induced cardiovascular responses predict post-sleep cardiac functions

We then established how these induced cardiovascular responses during sleep slow waves predict post-sleep left-ventricular systolic and diastolic function changes that we have previously reported in ^10^. Therefore, we investigated how the strength of the induced cardiovascular responses is related to post-sleep cardiac parameters. We extracted the cardiovascular responses within a window spanning from 2 seconds before to 18 seconds after the detected onset of both types of slow waves, that occurred during marked stimulation windows of all conditions (note that for the SHAM night, slow waves were only detected but not stimulated). The average of each cardiovascular response for every subject across each night was computed, followed by calculating the integrated area under the curve (iAUC) to ensure a positive valuation of each area under the response curve. Afterwards, we used repeated measures correlations between the iAUC cardiovascular metrics (instantaneous heart rate (IHR), systolic blood pressure (SBP), and diastolic blood pressure (DBP)) during sleep and post-sleep TTE-derived parameters that reflect cardiac function (global longitudinal strain, left-ventricular ejection fraction (LVEF), ratio of early diastolic filling to annular velocity (E/e’), and tricuspid annular plane systolic excursion (TAPSE)). As illustrated in Figure 3, we found a strong negative repeated measures correlation between the strength of IHR, SBP, DBP, and global longitudinal strain. Global longitudinal strain maps the complex contraction of the left-ventricular myocardium, predominately of the subendocardial longitudinally oriented fibers ^21^, and is an established marker assessing left-ventricular systolic function ^22,23^. Our results indicate increased left-ventricular systolic function (evidenced by a more negative global longitudinal strain) with an increase in cardiovascular response. Furthermore, we found strong positive repeated measures correlations between the cardiovascular responses and LVEF, the current standard clinical method to assess left-ventricular systolic function (see Table 1) ^24^. E/e’ was negatively correlated with the peak-to-peak IHR and blood pressure response (see Table 1), indicating increased left-ventricular diastolic function ^25^. Therefore, pointing towards increased relaxation of the left cardiac ventricle relating to increased cardiovascular response. Lastly, we investigated whether the cardiovascular iAUC metrics influence right-ventricular systolic function and, therefore, the contraction of the right ventricle (TAPSE). This cardiac metric was not reported previously in ^10^. Using a linear-mixed effects model approach, we found stimulation conditions to significantly influence post-sleep TAPSE (F(2,34)=4.7, p=0.016, see supplementary Figure 1), and post-hoc comparisons using paired t-tests revealing a significant increase in TAPSE (t(17) = 2.78, p=0.03) for *ISI*1_High_ compared to SHAM, whereas there was no significant difference between *ISI*1_Low_ and SHAM (t(17) = 1.47, p = 0.32)). Therefore, *ISI1*_High_ significantly increased post-sleep right-ventricular systolic function compared to SHAM. Investigating whether the cardiovascular iAUC response can predict these changes in TAPSE, we found a significant positive linear repeated measures correlation between the induced cardiovascular response and TAPSE (see Table 1), indicating increased right-ventricular systolic function with increased cardiovascular response.

**Table 1.** Repeated measures correlation analysis between post-sleep echocardiography parameters and cardiovascular responses after detected slow waves. To quantify cardiovascular responses to slow waves we used the integrated area under the curve for the echocardiographic parameters left-ventricular ejection fraction (LVEF), and tricuspid annular plane systolic excursion (TAPSE), or peak-to-peak distance for the E/e’ ratio. The analysis involved 15 subjects for instantaneous heart rate (IHR) and 17 subjects over three nights for diastolic (DBP) and systolic blood pressure (SBP).

|  | IHR |  | SBP |  | DBP |  |
| --- | --- | --- | --- | --- | --- | --- |
|  | p-value | r | p-value | r | p-value | r |
| LVEF | $p < 0.01$ | 0.46 | $p < 0.001$ | 0.56 | $p < 0.001$ | 0.53 |
| E/e' | $p < 0.1$ | -0.45 | $p < 0.1$ | -0.29 | $p < 0.1$ | -0.35 |
| TAPSE | $p < 0.001$ | 0.50 | $p < 0.001$ | 0.45 | $p < 0.001$ | 0.49 |

**Fig. 3.**
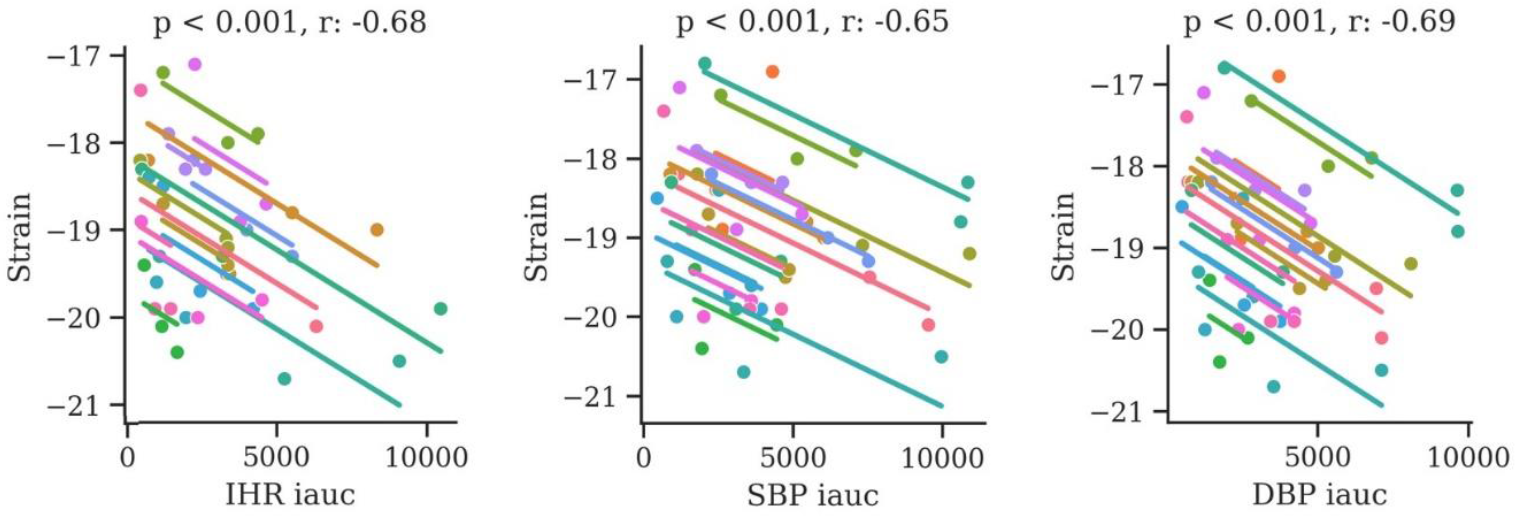
Repeated measures correlation of post-sleep global longitudinal strain with cardiovascular responses to detected slow waves in auditory stimulation windows. Data is incorporated for 15 subjects for instantaneous heart rate (IHR). and 17 subjects for diastolic and systolic blood pressure (DBP, SBP). Cardiovascular metrics were computed as integrated area under the curve (iAUC) and quantified in a time frame extending from 2 seconds before to 18 seconds after each slow wave onset occurring in a simulation window.

### Type I and type II slow waves are linked to different heart rate and blood pressure responses during unstimulated sleep

In the following step, our aim was to gain a deeper understanding of whether distinct slow waves are also occurring alongside distinct cardiovascular responses. Few studies proposed that strongly synchronized slow waves, such as K-complexes, evoke a cardiovascular activation response ^15,18^. Therefore, we classified the detected slow waves into type I and type II (see Methods) during periods of unstimulated sleep (SHAM condition) and investigated the cardiovascular responses associated with each type of slow wave. The cardiovascular responses occurring within the 18 seconds following the onset of each slow wave were normalized by dividing by the values from the 2 seconds preceding each onset. Subsequently, we averaged these normalized cardiovascular responses for each type of slow wave within each subject, and then across all participants. As illustrated in Figure 4, the IHR exhibited a rapid, almost symmetrical increase and decrease in response. In contrast, type II slow waves elicited a relatively weaker response across all cardiovascular metrics. In addition, our finding indicated a more pronounced cardiovascular blood pressure response with type I slow waves, peaking approximately 5 seconds after the wave onset. There were statistically significant differences (p<0.05) between type I and II slow waves in all cardiovascular metrics.

**Fig. 4.**
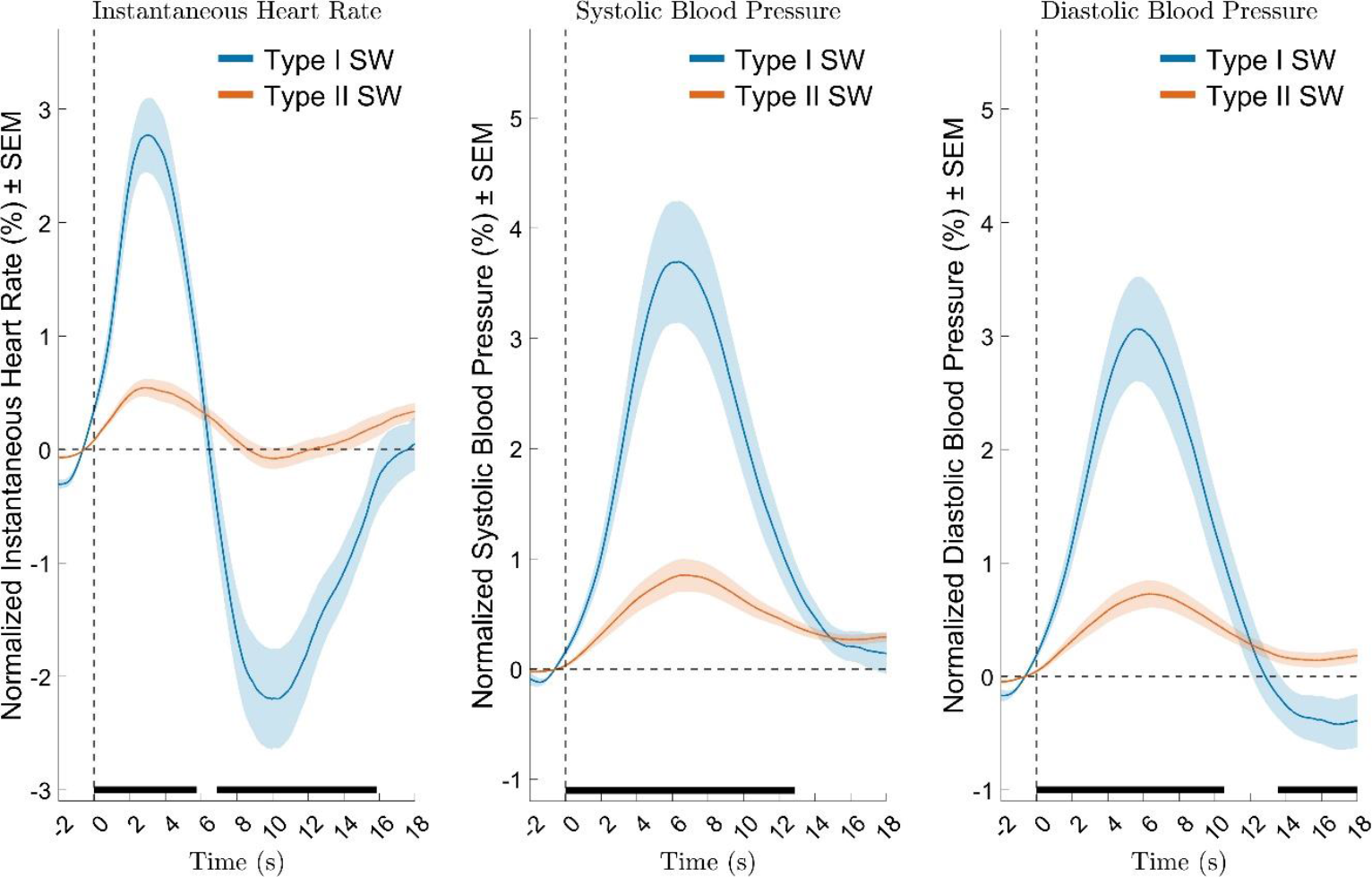
Distinct cardiovascular responses to type I and II slow waves in unstimulated sleep. Type I (including K-complexes) and type II slow waves show distinguishable responses, as evidenced in the data from 17 subjects for blood pressure and 14 for instantaneous heart rate across the unstimulated (SHAM) night. Stronger reactions in instantaneous heart rate, diastolic, and systolic blood pressure coincide with the onset (Time: 0s) of type I slow waves, whereas type II slow waves reveal smaller responses. The black line indicates significant differences (p<0.05) after slow wave onset between the slow wave types.

### Auditory stimulation increases the occurrence of type I and type II slow waves with distinct temporal dynamics

Because a strengthened cardiovascular response during sleep predicts post-sleep cardiac function (see Figure 3 and Table 1) we wanted to investigate whether auditory slow wave stimulation induces both types of slow waves similarly. Therefore, we identified occurrences of either type I or type II slow waves during the stimulation windows of all nights to calculate the likelihood of both types of slow waves occurring as a function of time during the stimulation window. The likelihood of each type was calculated by summing each of their occurrences, multiplying by 100, and then dividing these values against the total number of stimulation windows.

Figure 5 reveals that type I and type II slow wave likelihood differs in response to auditory stimulation. In the ON window, type II slow waves demonstrated an increase in wave likelihood in response to each tone played as stimulation was delivered in a rhythmic 1-Hz manner. Notably, the initial tones induced on average a higher type II wave likelihood than the later tones, suggesting that the first tones are more effective than the last ones in evoking a slow wave. Of note, the first decreasing drop at the start of the stimulation windows in type II slow waves is associated with the up-phase stimulation of ongoing slow waves (see Figure 5). For type I slow waves we observed a pronounced increase in wave likelihood to auditory stimulation primarily at the onset of the stimulation window compared to the SHAM condition. Type I likelihood is not displaying a rhythmic response as observed for type II slow waves. Furthermore, the occurrence of type I slow waves is lower compared to type II slow waves.

**Fig. 5.**
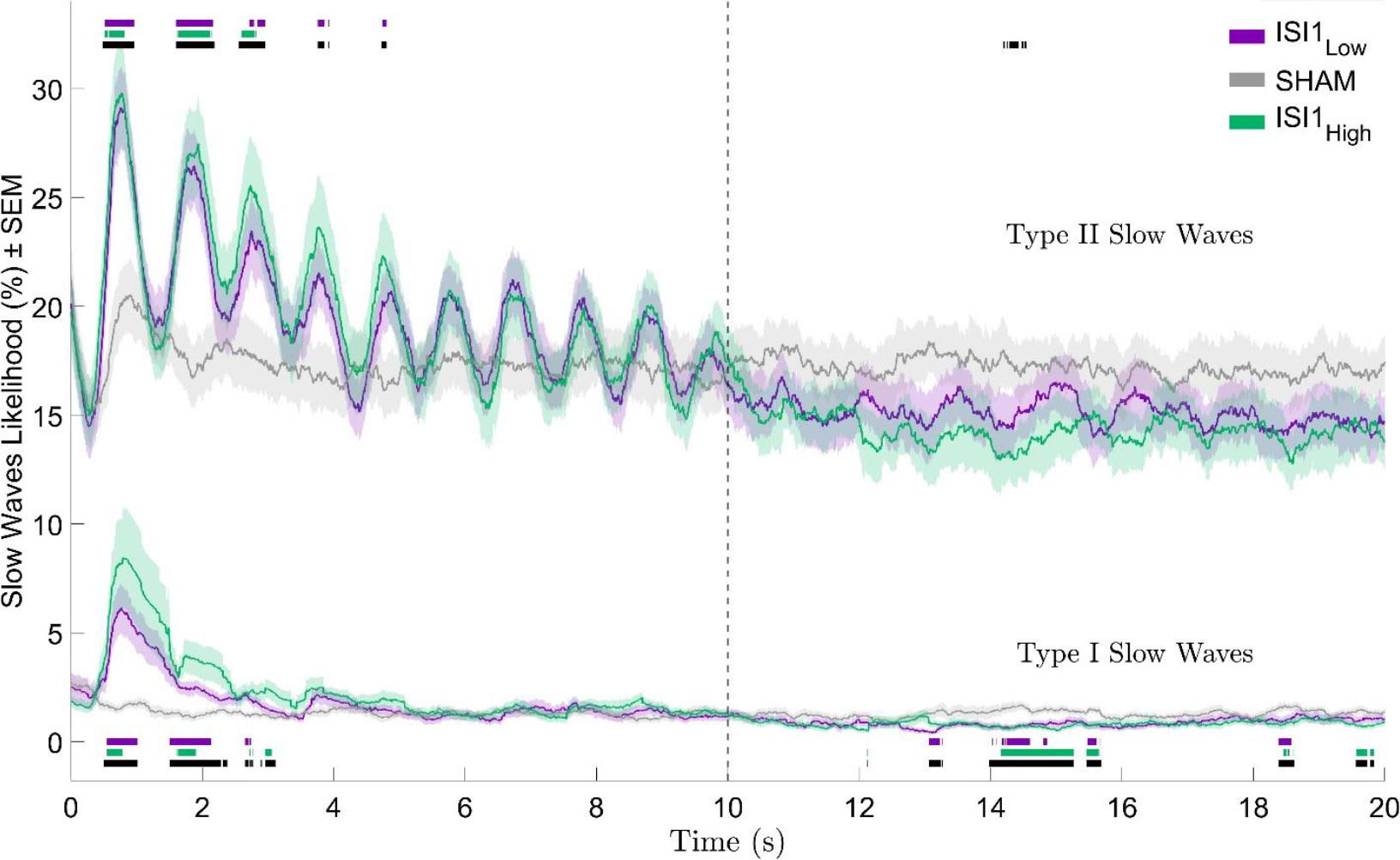
Different impact of auditory stimulation on type I and type II slow waves. Auditory stimulation differentially affects type I (including K-complexes) and type II slow waves occurrences, based on data from 17 subjects over three experimental night conditions. Type II slow waves show an increase in likelihood during ON window stimulation each second, followed by a rapid decrease, a trend consistent across different volume levels. In contrast, type I slow waves (including K-complexes), are characterized by a pronounced initial response. Black lines indicate significant (p<0.05) difference between all conditions, whereas purple and green lines indicate significant post-hoc comparisons between auditory stimulation conditions and SHAM control with *ISI1*_Low_ and *ISI1*_High_, respectively.

## Discussion

Our work uncovers that post-sleep cardiac function is predicted by the strength of cardiovascular responses induced alongside slow waves. First, we confirmed our previous work ^10,12^ showing that auditory slow wave stimulation evokes a cardiovascular activation response, resembling cardiovascular pulsations. We then demonstrate that the strength of these cardiovascular pulsations is correlated with TTE-derived cardiac left-ventricular and right-ventricular function the next morning. We further found that type I and type II slow waves are not only functionally distinct based on EEG-based wave morphology, timing, and synchronization processes ^13,14^, but also signify unique brain and body “micro-states” during sleep that may be differently involved in promoting cardiac function. Specifically, both slow wave types are associated with distinct strengths of cardiovascular pulsations. Therefore, we provide mechanistic insights into how these two slow wave types are related to post-sleep cardiac function. Lastly, we showed that auditory slow wave stimulation enhances the likelihood of both, type I and II slow waves compared to the SHAM control condition, however, with different underlying temporal dynamics. Therefore, altogether, we show a pathway on how auditory slow wave stimulation may promote cardiac function during sleep.

### Type I slow waves align with strengthened cortico-cardiovascular coupling, predicting improved cardiovascular function

We here describe a mechanistic explanation of how slow waves may increase cardiac function. To date, there is only limited evidence highlighting the importance of deep NREM sleep and slow waves in promoting cardiovascular health ^9,15,26,27^. We demonstrate that the strength of cardiovascular pulsations induced alongside slow waves is correlated to enhancements in cardiac functions after sleep (see Figure 3 and Table 1). More specifically, the average cortico-cardiovascular coupling during sleep predicts these cardiac improvements. Additionally, this cortico-cardiovascular coupling is stronger in type I slow waves compared to type II slow waves (see Figure 4), and therefore, type I slow waves potentially have stronger implications on post-sleep cardiovascular functions than type II slow waves.

We furthermore show a cardiovascular activation as a response with type I slow waves, which is similar to the response previously reported cardiovascular with K-complexes ^15,17,18^. Additionally, De Zambotti et al. ^15^ demonstrated that played tones without eliciting a K-complex did not evoke a biphasic cardiac response. They found that both naturally occurring and tone-evoked K-complexes elicit similar cardiac responses. Hence, an observable K-complex in the sleep EEG serves as a necessary predictor or initiator for a cardiovascular response to manifest. Nevertheless, our study confirms the biphasic heart rate response observed in both stimulated and spontaneous type I slow waves comparable to K-complexes ^15,26^. Therefore, our findings support the hypothesis of an involvement of K-complexes or type I slow waves in blood pressure ^18^ and cardiac ^15^ regulation, possibly directly influencing cardiovascular function during sleep ^15,17,18,26^. Thus, we provide evidence that the mechanisms underlying the generation of type I slow waves are also modulating the cardiovascular system during sleep ^15^, highlighting their importance as a fundamental sleep element associated with promoting cardiovascular health ^10^. Moreover, our findings propose a potential mechanistic explanation for aging as the major risk factor for cardiovascular diseases ^28^, because aging is associated with a strong decline in K-complex density ^29^. Altogether, auditory slow wave stimulation protocols aiming to increase cardiovascular function should focus on increasing type I slow wave occurrence and essentially its associated cardiovascular pulsations.

### Type I and type II slow waves signify unique brain and body microstates

We observed coupled cardiovascular pulsations being evoked in association with both types of slow waves. The increase in response in type I slow waves could be caused by the stronger activation of brain structures that are also involved in autonomic modulation of the cardiovascular system. On an EEG morphological level, the two distinct types of slow waves have been proposed to be generated from distinct generation and synchronization mechanisms within the central nervous system ^13^. More specifically, type I slow waves have been suggested to manifest through subcortico-cortical synchronization processes, whereas type II slow waves emerge through cortico-cortical synchronization ^13,14^. Our observed stronger cortico-cardiovascular coupling in type I slow waves supports the idea of an enhanced subcortical involvement because the processes underlying type I slow waves likely activate the sympathoexcitatory locus coeruleus ^30^, a small nucleus in the brain stem serving as noradrenergic center of the brain. Supporting this idea, we recently showed a pupil dilation alongside K-complexes ^31^, indirectly pointing towards a potential increase in locus coeruleus activity concurrent with highly-synchronized slow waves ^32^. Additionally, K-complexes coincide with spikes in muscle sympathetic nerve activity ^18,20^, also supporting the stronger cardiovascular activation response compared to type II slow waves. Altogether, type I and type II slow waves are not only distinguishable by cortical characteristics, but also by their different cardiovascular responses.

### Slow wave stimulation as a therapeutic tool to enhance cardiovascular health

Our findings emphasize the potential of slow wave stimulation to strengthen cortico-cardiovascular coupling as a tool for improved cardiovascular function. We observed a significant correlation between the cortico-cardiovascular coupling and two markers estimating left-ventricular systolic function. On the one hand, strain maps the complex contraction of the left-ventricular myocardium ^21^, and has been established as an important marker indicating deformation of the left ventricle and for assessing left-ventricular systolic function ^22,23^. On the other hand, LVEF, the current standard clinical method to assess left-ventricular systolic function represents an important parameter for the primary and secondary prevention of cardiac diseases ^33,34^. In addition to systolic function, we also foundasignificant correlation between the cortico-cardiovascular coupling and decreased E/e’ ratio. The left-ventricular diastolic function reflects relaxation and stiffness of the myocardium ^25,35^. Particularly, an E/e’ ratio>14 has been related to abnormal diastolic function of the heart ^25^. In addition to left-ventricular function, we also found a positive correlation between the cardiovascular response during sleep TAPSE. TAPSE has been associated with increased risks for right-ventricular myocardial infarction ^36^, and is proposed to be an important predictor for survival in cardiopulmonary diseases ^37^. Therefore, promoting such cardiovascular pulsations in relation to induced and natural slow waves during sleep could be another important preventative pathway in maintaining cardiovascular health. Given that this cortico-cardiovascular coupling is enhanced in type I slow waves compared to type II slow waves, future stimulation protocols should aim to enhance type I slow waves or K-complexes.

Most K-complexes occur spontaneously during NREM sleep, without any external cause ^19,38,39^. However, K-complexes can also be evoked through somatosensory stimulation such as auditory stimulation ^30^. We show here that both, type I and type II slow waves can be enhanced through auditory stimulation (see Figure 5). Type I slow waves were evoked only at the start of each stimulation window, potentially because of a refractory period as has been reported for K-complexes ^13,40,41^. Type II slow waves, on the other hand, can be evoked or even entrained with each single presented tone. Moreover, we also confirm our previous findings ^10,12^ that auditory stimulation strengthens cardiovascular pulsations compared to the SHAM control condition. These findings are in line with early studies reporting an autonomic activation response observable after scored K-complexes ^18,42^. This activation response could even be attributed to dampening the baroreflex, leading to sleep continuity ^18^. However, both types of slow waves likely contribute to cardiac fluctuations and therefore, may even increase heart rate variability ^15^, but to a different extent. However, in order to improve cardiovascular health, we propose to develop auditory stimulation protocols to optimize the occurrence of these cardiovascular fluctuations and strength of cardiovascular pulsations, by maximizing type I slow wave occurrence.

Altogether, our findings underscore the potential of sleep slow wave stimulation as a therapeutic tool to enhance cardiovascular function. By differentiating the responses of type I and II slow waves, our study contributes to a deeper understanding of how slow waves may be strategically influenced to optimize cardiovascular health by strengthening cortico-cardiovascularcoupling. This advancement holds promise for developing targeted auditory stimulation strategies, potentially offering a non-invasive means to enhance cardiovascular recovery during sleep.

### Cortico-cardiovascular coupling likely deteriorates in neurodegenerative disorders

Besides cardiovascular diseases, Alzheimer’s disease represents another aging disease with shared risk factors that are associated with changes in sleep ^43–45^. Recent studies underscore the crucial role of sleep, particularly NREM sleep, in facilitating the removal of metabolic waste from the brain through the glymphatic system ^46,47^. Mainly with a focus on improving memory functions, auditory stimulation during sleep has been employed to increase slow waves ^48,49^, and has been related to more favorable amyloid plasma response. Recently it has been shown that plasma proteomic signatures indicating accelerated heart aging predict Alzheimer’s disease progression ^50^, again emphasizing a tight link between sleep, cardiovascular function, and neurodegenerative disorders such as Alzheimer’s disease. Interestingly, the process of metabolic brain clearance seems to be driven by vasomotor pulsations throughout the brain ^51^, which are potentially directed by autonomic arousals ^52^. These pulsations, in turn, amplify the flow of cerebrospinal fluid ^53^. The here described and observed cortico-cardiovascular coupling is consistent with the reported autonomic arousals ^52^ to cleanse the brain. Therefore, the decrease in strength of this coupling could represent an important mechanism of how the degenerating brain in Alzheimer’s disease pre-clinical stage results in the accumulation of amyloid plaque. Essentially, increasing this cortico-cardiovascular coupling during sleep by evoking type I slow waves or K-complexes might increase cerebrospinal fluid flow and thereby, decelerate amyloid accumulation.

### Limitations

Our study only investigated healthy male participants between 30 and 60 years of age. The female menstrual cycle has been shown to influence sleep ^54^ and also HR during sleep ^55^. Moreover, menopause generally occurs in women within our targeted age range, coinciding with sleep modifications and disturbances ^56^. These hormonal differences would make it difficult to investigate the effects of slow waves on cardiovascular functions across sessions and to generalize findings across participants. Therefore, for this initial study, we decided to only include male participants. Nevertheless, future large-scale studies should also explore type I and type II slow waves and their relationship with cardiovascular function across sex. Moreover, the number of participants in this study appears relatively low. While this number is comparable to other interventional sleep studies ^6–8^,a larger scale study applying auditory stimulation across multiple nights and in a broader and more heterogenous population sample is needed to characterize long-term effects. In this study, we tested healthy participants. However, to verify the prospective clinical application for enhancing cardiac function in individuals with heightened cardiovascular risks or existing cardiovascular conditions, it is essential to first establish the preventive and therapeutic effects associated with increased cortico-cardiovascular coupling during sleep.

In this study, we present insights into the cardiovascular implications of slow waves during sleep. Our findings reveal that auditory slow wave stimulation strengthens cortico-cardiovascular coupling, and its strength predicts cardiac function. Particularly type I slow waves, including K-complexes, co-occur with stronger cardiovascular pulsations compared to type II slow waves. Therefore, this indicates that both types of slow waves represent unique brain and body micro-states. Moreover, we showed that auditory stimulation differently intensifies both types of slow waves. By establishing the connection between slow wave types and their cardiovascular responses, our study marks a strong advancement in decoding the complex relationship between sleep physiology and cardiovascular health.

## Materials and Methods

### participants and experimental procedure

Overall, 29 healthy male participants were enrolled in the randomized-controlled trial (registered at ClinicalTrials.gov (NCT04166916)) . The here presented analysis is a secondary analysis of the results reported previously ^10^. Participants were recruited from the local community using advertisements on different platforms. All participants were of good general health, non-smokers, reported a regular sleep-wake rhythm, had a body mass index between 17-30, and none of the participants had any cardiovascular disorder, which has been confirmed by an expert cardiologist. Furthermore, all participants had no psychiatric or neurological disease and no presence or suspicion of any sleep disorders. Exclusion criteria further encompassed any participant using on-label sleep medication or medication impacting cardiovascular function, as well as those demonstrating low sleep quality during the screening night (defined as sleep efficiency below 75%). Finally, 18 participants underwent the full three nights auditory stimulation protocol. All included participants provided written informed consent prior to the study and were compensated for their participation. The study was conducted following the declaration of Helsinki and approved by the cantonal Ethics Committee Zurich (BASEC2019-01538).

All participants underwent a screening period including a phone screening and a screening night to familiarize participants with the sleep lab environment and all measurement devices. To ensure hearing capabilities of 40 dB, we conducted a short audiometry test. After participants had passed the screening night, they were screened by an expert cardiologist for any cardiovascular comorbidities at the University Hospital Zurich before the first experimental night. Participants were asked to adhere to a regular sleep rhythm one week before the start of the experimental night (sleep duration of 7-9 hours, bedtime *±* 1 hour to the scheduled bedtime in the sleep lab, no naps). Three nights before the main experimental night, participants had to sleep for 7.5 hours *±* 0.5 hours, with a bedtime *±* 0.5 hour to the scheduled bedtime. Adherence was verified through actigraphy (GENEActiv, Activinsights, Kimbolton, UK) and a sleep diary.

On the day of each experimental night, participants had to abstain from naps, alcohol, intensive sports, caffeine, and food 5 hours before arriving at the sleep lab. We attached a high-density EEG system (Geodesic Sensor Net, Magstim EGI, Eugene, USA), and two ECG electrodes (Gold Genuine Grass electrodes, Natus Medical Inc., Pleasanton, US) under the right collar bone and on the left rib-cage, respectively. All impedances of EEG and ECG electrodes were kept below 40 k ohm. Afterward, a continuous non-invasive blood pressure monitoring system (Finapres Novascope, Finapres Medical Systems BV, Enschede, Netherlands) was attached and calibrated to ensure proper attachment and signal quality. After lights out, participants could sleep for 7.5 hours while polysomnography and ECG were recorded (BrainProducts GmbH, Gliching, Germany) at a sampling rate of 500 Hz. Beat-to-beat blood pressure was continuously measured with a sampling rate of 200 Hz. After a constant morning routine including additional tests (including autonomic maneuvers, and cognitive and behavioral testing, not reported here), participants went to the University Hospital Zurich for a TTE measurement.

In our study, data integrity was crucial in determining the number of participants included in specific analyses. Because of a technical failure in acquiring the overnight EEG data of the SHAM night of one participant, this participant was completely excluded from all analyses, resulting in 17 subjects included in the here presented analyses (all males, aged 45.5 ± 9.86 years). Challenges encountered during ECG data acquisition compromised the reliability of specific measurements, necessitating a reduction in the participant count to 14 or 15. Specifically, for the analyses leading to Figures 2 and 3, we had no choice but to exclude two participants regarding the generation of the IHR plot. Their ECG signals were compromised specifically during the critical periods of stimulated slow-wave activity we aimed to analyze. Segments from the ECG signal were considered compromised whether R peaks were not detected due to poor quality data. Furthermore, the analysis presented in Figure 4 not only targeted stimulated slow waves but also categorized them by type (I or II). This additional stratification, beyond what was necessary for Figures 2 and 3, led us to exclude an additional participant due to a combination of ECG signal corruption and a low incidence of type I stimulated slow waves. Consequently, the IHR plot in Figure 4 was derived from data from 14 subjects, reflecting the stringent criteria applied to ensure data integrity.

### Auditory slow wave stimulation protocol

To investigate the effects of slow waves on cardiovascular function, participants completed a randomized crossover experimental protocol with three stimulation nights. We applied two stimulation conditions that were selected based on their dynamic effects on the brain and cardiovascular system ^10^ and compared them to a SHAM (no stimulation) condition. Namely, we used 1 Hz rhythmic EEG feedback-controlled auditory stimulation of a 50 ms burst of pink noise with a sound level of 45 dB (*ISI*1_High_) and a sound level of 42.5dB (*ISI*1_Low_). We used two different sound volumes to achieve a dose-dependency effect and to investigate which dose is still effective in inducing possible cardiovascular responses. Auditory stimulation was delivered using a real-time system programmed in OpenViBE ^57^, described in more detail in previous work ^10^. Whenever all stimulation conditions to ensure stimulation during NREM sleep were met, auditory stimulations were presented through Etymotic insert earphones (Etymotic Research Inc., ER 3C). Stimulations were delivered in a windowed ten-second ON (auditory stimulation presented) followed by a ten-second OFF (no auditory stimulation presented) design. The first stimulation of each ON stimulation window targeted the up-phase of a detected slow wave. Four hours after the first stimulation window started, stimulation was halted for the remainder of the night.

### EEG data processing and slow wave classification

Data processing was conducted in MATLAB (R2022b, MathWorks Inc., Natick, MA), with the initial EEG recording, sampled at 500 Hz, being down-sampled to 200 Hz. Employing the EEGLAB toolbox ^58^, pre-processing involved detrending to remove slow drifts and low-frequency artifacts, using a 1.5-second sliding window with a 0.02 seconds step. The PREP pipeline algorithm ^59^ was used to identify poor-quality EEG channels through amplitude assessment, signal-to-noise ratio estimation, inter-channel correlation analysis, and RANSAC-based channel prediction. Identified noisy channels were temporarily excluded to prevent data distortion. Further, we applied a 50 Hz notch filter and a 0.5-45 Hz Butterworth bandpass filter using TESA within EEGLAB ^60^. Independent Component Analysis was then conducted to decompose EEG recordings, isolating neural from non-neural signal ^61–63^. Components were classified using the *ICLabel* algorithm ^64^, distinguishing between brain-related and non-brain-related activities. A rigorous exclusion criterion based on confidence levels and comparative ratios was applied to retain only relevant neural components. Finally, previously excluded noisy channels were reintegrated using spherical spline interpolation and the data was re-referenced to mastoid electrodes for further analysis.

An automated, two-step algorithm was employed to identify slow waves in EEG recordings ^13^. The first step involved reducing multi-channel EEG data to a single-channel representation. For each time-point, we selected and averaged the three most negative amplitude values across all available channels after excluding the most negative extreme values and data from peripherally situated electrodes. This procedure allowed us to obtain a composite signal representing the negative envelope of the EEG signals across the considered set of electrodes. A Butterworth band-pass filter with a frequency range of 0.5 to 40 Hz was applied to obtain both zero-mean signal centering and noise reduction. Potential slow waves were then selected using a duration-based criterion applied to detected negative half-waves. Specifically, a waveform was classified as a slow wave if its negative component had a duration between 0.25 and 1 second and it was observed during the NREM 2 or NREM 3 sleep stages.

Following the identification of slow waves in EEG recordings, we conducted a detailed examination of individual slow wave events, extracting their morphological and dynamic features. These features included the negative slope (*μ*V*/*s), calculated as the gradient between the wave’s onset and its maximum negative peak, reflecting the rate of neuronal down-state entry ^65,66^. The positive slope (*μ*V*/*s), computed from the wave’s maximum negative peak to the point where the waveform becomes positive, quantified the transition into an up-state ^65,66^. Additionally, an involvement score (dimensionless) was determined, representing the percentage of channels contributing to the slow wave, calculated around the maximum negative peak with a fixed amplitude threshold (-5*μ*V).

These features led to the definition of the synchronization score (mV*/*s), a key metric for distinguishing slow wave types based on cortical involvement and synchronization speed ^65^. The synchronization score was instrumental in classifying slow waves into type I and type II categories, based on dynamic thresholds derived from the synchronization score distribution in each EEG recording. After calculating the synchronization scores for each EEG recording, two critical thresholds were determined to categorize slow waves. These thresholds were calculated by adding the median value of the synchronization scores to its median absolute error, multiplied by a specific numerical factor. A factor of 3 defined the lower threshold, while a factor of 4 established the upper threshold. Consequently, slow waves with synchronization scores below the lower threshold were classified as Type II, and those above the upper threshold as Type I. Slow waves falling between these two thresholds were marked as indeterminate and excluded from analysis when focusing on the differences between types I and II.

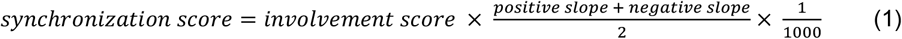

Finally, an additional classification layer was implemented to distinguish stimulated slow waves from spontaneous ones, depending on whether they coincided with the ON period of the stimulation windows. It is crucial to note that during the SHAM condition, stimulated slow waves are, spontaneous slow waves occurring within the ON windows, which are automatically set through the EEG-feedback algorithm, serving as a control comparison.

### Heart rate and blood pressure during sleep

Continuous blood pressure and heart rate are derived 2 seconds before to 18 seconds after each slow wave onset, respectively from the continuous non-invasive blood pressure monitoring system and the electrocardiogram. These data, extracted within the stimulation windows, were reported here ^10^. Briefly, electrocardiogram R-waves were detected automatically and visually corrected if necessary using PhysioZoo ^67^. Segments of non-detectable R-waves due to poor data quality were excluded. The generated RR interval sequences were further processed with MATLAB. IHR was calculated by dividing 60 through the specific RR interval and interpolating the beat-to-beat data to 200 Hz by using linear interpolation. Segments of bad quality or artifacts of nocturnal diastolic and systolic blood pressure were removed based on detected heart rate outliers, using the function *isoutlier* with the option *gesd* in MATLAB. We further removed segments with less than three valid data points within ten seconds, as the low number of valid measurements could indicate bad recording quality. Blood pressure data points were interpolated to 200 Hz by using linear interpolation.

### Echocardiography

Each morning following the experimental night, an experienced cardiologist performed a TTE on a GE Healthcare E95 echocardiography station (GE Healthcare Systems, Wisconsin, USA). The following measurements were collected: Left-ventricular end-diastolic diameter (LVEDD), left-ventricular end-diastolic volume (LVEDV), left ventricular end-systolic volume (LVESV), LVEF, left-ventricular longitudinal strain (strain), left-ventricular tissue doppler (E-wave, A-wave, E/A ratio, lateral e’ wave, E/e’ ratio), trans-tricuspidal systolic right ventricular gradient, right ventricular end-diastolic area (RVareaD), and TAPSE. The findings related to auditory stimulation conditions were partially detailed here ^10^.

### Statistical analysis

Statistical analyses were conducted in Python (version 3.9.13), MATLAB (version R2022b), and R ^68^ (R Studio version 1.2.5033). To compare post-sleep TTE parameters between conditions that have not been reported in ^10^, we used linear mixed-effects models with fixed factors condition and random factor subject using the software package lme ^69^ in R. Here, if the model was significant for condition, we derived post-hoc p-values for both auditory stimulation conditions compared to SHAM using paired t-tests and correcting for multiple comparisons applying the Bonferroni correction. Using Python, we computed the repeated measures linear correlation ^70^ to correlate TTE parameters with cardiovascular responses from auditory-stimulated slow waves. For Figures 2, 4, and 5, we performed linear-mixed effects models using the MATLAB function *fitmod*, entering subject as random factor and condition as fixed factor (*α*=0.05) for every timepoint. Post-hoc p-values were obtained through paired t-tests between ISI1_High_ and SHAM, and *ISI1*_Low_ and SHAM. Given the repeated measures analysis for each timepoint, we applied the false discovery rate Benjamini-Hochberg correction for multiple comparisons for the p-values obtained through the linear-mixed effects models and the post-hoc testing. Post-hoc p-values were in addition corrected for multiple comparisons between conditions by using the Bonferroni-corrected p-value threshold of 0.025. Otherwise, p < 0.05 was considered as significant.

## Acknowledgments

All authors thank their trainees, collaborators, and mentors for the expiring exchanges and dialogues on the thematic of this article. SleepLoop consortium members provided uncountable discussions and feedback. We are grateful to the Alumni Association of ETH Zurich for their support in participant recruitment for this study. We also thank Dr. Rafael Polanía for his continuous support and feedback during the study.

## Funding

Swiss National Science Foundation PZ00P3_179795

SleepLoop Flagship of Hochschulmedizin Zürich

Swiss National Science Foundation Grant 32003B_207719

National Research Foundation, Prime Minister’s Office, Singapore under its Campus for Research

Excellence and Technological Enterprise (CREATE) program (FHT)

Swiss National Science Foundation 320030_179443

Italian Ministry of University and Research - PON Program Green and Innovation, 2014-2020

European Research Council (ERC) Starting Grant 948891

## Author contributions

Conceptualization: GA, SH, RH, NW, CS, CL

Methodology: GA, GB (Type I & Type II); SH, CL (auditory stimulation)

Software: GA, FSi, GB (Type I & Type II); SH (auditory stimulation)

Validation: GA

Formal analysis: GA, SH

Investigation: SH, MCD, FS, RS, FA, AT, SM, DN, PB, CS

Resources: NW, CS, CL

Data Curation: GA, SH

Visualization: GA, SH

Supervision: NW, CL

Writing—original draft: GA, SH, CL

Writing—review and editing: GA, SH, GB, MCD, FS, RS, FA, AT, SM, DN, PB, PC, FSi, RH, NW, CS, CL

Project administration: SH, CL

Funding acquisition: NW, PC, CL

## References

1. Valenzuela P, Santos-Lozano A, Torres-Barrán A, Morales JS, Castillo-García A, Ruilope L, Insua DR, Ordovás J, Lucia A. Poor self-reported sleep is associated with risk factors for cardiovascular disease: A cross-sectional analysis in half a million adults. Eur J Clin Invest 2021;52.

2. Kohansieh M, Makaryus AN. Sleep Deficiency and Deprivation Leading to Cardiovascular Disease. Int J Hypertens 2015;2015:1–5.

3. Cappuccio FP, Cooper D, D’Elia L, Strazzullo P, Miller MA. Sleep duration predicts cardiovascular outcomes: a systematic review and meta-analysis of prospective studies. Eur Heart J 2011;32:1484–1492.

4. Grandner MA, Hale L, Moore M, Patel NP. Mortality associated with short sleep duration: the evidence, the possible mechanisms, and the future. Sleep Med Rev 2010;14:191–203.

5. Lustenberger C, Ferster ML, Huwiler S, Brogli L, Werth E, Huber R, Karlen W. Auditory deep sleep stimulation in older adults at home: a randomized crossover trial. Commun Med 2022;2:30.

6. Ngo H-V V., Martinetz T, Born J, Mölle M, Molle M. Auditory Closed-Loop Stimulation of the Sleep Slow Oscillation Enhances Memory. Neuron 2013;78:545–553.

7. Ngo H-V V, Miedema A, Faude I, Martinetz T, Mölle M, Born J. Driving Sleep Slow Oscillations by Auditory Closed-Loop Stimulation—A Self-Limiting Process. J Neurosci 2015;35:6630–6638.

8. Besedovsky L, Ngo H-V V., Dimitrov S, Gassenmaier C, Lehmann R, Born J. Auditory closed-loop stimulation of EEG slow oscillations strengthens sleep and signs of its immune-supportive function. Nat Commun 2017;8:1984.

9. Grimaldi D, Papalambros NA, Reid KJ, Abbott SM, Malkani RG, Gendy M, Iwanaszko M, Braun RI, Sanchez DJ, Paller KA, Zee PC. Strengthening sleep-autonomic interaction via acoustic enhancement of slow oscillations. Sleep 2019;42:1–11.

10. Huwiler S, Carro-Domínguez M, Stich FM, Sala R, Aziri F, Trippel A, Ryf T, Markendorf S, Niederseer D, Bohm P, Stoll G, Laubscher L, Thevan J, Spengler CM, Gawinecka J, Osto E, Huber R, Wenderoth N, Schmied C, Lustenberger C, Carro-Dominguez M, Stich FM, Sala R, Aziri F, Trippel A, Ryf T, Markendorf S, Niederseer D, Bohm P, Stoll G, Laubscher L, Thevan J, Spengler CM, Gawinecka J, Osto E, Huber R, Wenderoth N, Schmied C, Lustenberger C. Auditory stimulation of sleep slow waves enhances left ventricular function in humans. Eur Heart J 2023;44:4288–4291.

11. Diep C, Ftouni S, Drummond SPA, Garcia-Molina G, Anderson C. Heart rate variability increases following automated acoustic slow wave sleep enhancement. J Sleep Res 2022;31:1–6.

12. Huwiler S, Domínguez M, Huwyler S, Kiener L, Stich FM, Sala R, Aziri F, Trippel A, Schmied C, Huber R, Wenderoth N, Lustenberger C, Carro Dominguez M, Huwyler S, Kiener L, Stich FM, Sala R, Aziri F, Trippel A, Schmied C, Huber R, Wenderoth N, Lustenberger C. Effects of auditory sleep modulation approaches on brain oscillatory and cardiovascular dynamics. Sleep 2022;45:1–36.

13. Bernardi G, Siclari F, Handjaras G, Riedner BA, Tononi G. Local and widespread slow waves in stable NREM sleep: Evidence for distinct regulation mechanisms. Front Hum Neurosci 2018;12:1– 13.

14. Siclari F, LaRocque JJ, Postle BR, Tononi G, Bernardi G, Riedner BA, LaRocque JJ, Benca RM, Tononi G. Two distinct synchronization processes in the transition to sleep: A high-density electroencephalographic study. Sleep 2014;37:1621–1637F.

15. Zambotti M de, Willoughby AR, Franzen PL, Clark DB, Baker FC, Colrain IM. K-Complexes: Interaction between the Central and Autonomic Nervous Systems during Sleep. Sleep 2016;39:1129–1137.

16. Greenlund IM, Smoot CA, Carter JR. Sex differences in blood pressure responsiveness to spontaneous K-complexes during stage II sleep. 2021.

17. Sforza E, Jouny C, Ibanez V. Cardiac activation during arousal in humans: Further evidence for hierarchy in the arousal response. Clin Neurophysiol 2000;111:1611–1619.

18. Tank J, Diedrich A, Hale N, Niaz FE, Furlan R, Robertson RM, Mosqueda-Garcia R. Relationship between blood pressure, sleep K-complexes, and muscle sympathetic nerve activity in humans. Am J Physiol - Regul Integr Comp Physiol 2003;285:208–214.

19. Monstad P, Guilleminault C. Cardiovascular changes associated with spontaneous and evoked K-complexes. Neurosci Lett 1999;263:211–213.

20. Somers VK, Dyken ME, Mark AL, Abboud FM. Sympathetic-nerve activity during sleep in normal subjects. N Engl J Med 1993;328:303–307.

21. Sugimoto T, Dulgheru R, Bernard A, Ilardi F, Contu L, Addetia K, Caballero L, Akhaladze N, Athanassopoulos GD, Barone D, Baroni M, Cardim N, Hagendorff A, Hristova K, Lopez T, La Morena G De, Popescu BA, Moonen M, Penicka M, Ozyigit T, Carbonero JDR, Veire N Van De, Bardeleben RS Von, Vinereanu D, Zamorano JL, Go YY, Rosca M, Calin A, Magne J, Cosyns B, Marchetta S, Donal E, Habib G, Galderisi M, Badano LP, Lang RM, Lancellotti P. Echocardiographic reference ranges for normal left ventricular 2D strain: Results from the EACVI NORRE study. Eur Heart J Cardiovasc Imaging 2017;18:833–840.

22. Bergler-Klein J. Strain and left ventricular volumes for predicting cardiotoxicity: A life-saving approach in anthracycline cancer treatment? Eur Heart J Cardiovasc Imaging 2015;16:968–969.

23. McDonagh TA, Metra M, Adamo M, Gardner RS, Baumbach A, Böhm M, Burri H, Butler J, Celutkiene J, Chioncel O, Cleland JGF, Coats AJS, Crespo-Leiro MG, Farmakis D, Gilard M, Heymans S. 2021 ESC Guidelines for the diagnosis and treatment of acute and chronic heart failure. Eur Heart J 2021;42:3599–3726.

24. Klaeboe LG, Edvardsen T. Echocardiographic assessment of left ventricular systolic function. J Echocardiogr 2019;17:10–16.

25. Nagueh SF, Smiseth OA, Appleton CP, Byrd BF, Dokainish H, Edvardsen T, Flachskampf FA, Gillebert TC, Klein AL, Lancellotti P, Marino P, Oh JK, Popescu BA, Waggoner AD. Recommendations for the Evaluation of Left Ventricular Diastolic Function by Echocardiography: An Update from the American Society of Echocardiography and the European Association of Cardiovascular Imaging. J Am Soc Echocardiogr 2016;29:277–314.

26. Zambotti M de, Trinder J, Silvani A, Colrain IM, Baker FC. Dynamic coupling between the central and autonomic nervous systems during sleep: A review. Neurosci Biobehav Rev 2018;90:84–103.

27. Sayk F, Teckentrup C, Becker C, Heutling D, Wellhöner P, Lehnert H, Dodt C, Wellhoener P, Lehnert H, Dodt C. Effects of selective slow-wave sleep deprivation on nocturnal blood pressure dipping and daytime blood pressure regulation. Am J Physiol - Regul Integr Comp Physiol 2010;298:191–197.

28. North BJ, Sinclair DA. The Intersection Between Aging and Cardiovascular Disease. Circ Res 2012;110:1097–1108.

29. Crowley K, Trinder J, Kim Y, Carrington M, Colrain IM. The effects of normal aging on sleep spindle and K-complex production. Clin Neurophysiol 2002;113:1615–1622.

30. Bellesi M, Riedner BA, Garcia-Molina GN, Cirelli C, Tononi G. Enhancement of sleep slow waves: underlying mechanisms and practical consequences. Front Syst Neurosci 2014;8:82–88.

31. Carro-Domínguez M, Huwiler S, Oberlin S, Oesch TL, Badii G, Lüthi A, Wenderoth N, Meissner SN, Lustenberger C. Pupil size reveals arousal level dynamics in human sleep. 2023.

32. Murphy PR, O’Connell RG, O’Sullivan M, Robertson IH, Balsters JH. Pupil diameter covaries with BOLD activity in human locus coeruleus. Hum Brain Mapp 2014;35:4140–4154.

33. Wang TJ, Evans JC, Benjamin EJ, Levy D, LeRoy EC, Vasan RS. Natural history of asymptomatic left ventricular systolic dysfunction in the community. Circulation 2003;108:977–982.

34. Breathett K, Allen LA, Udelson J, Davis G, Bristow M. Changes in Left Ventricular Ejection Fraction Predict Survival and Hospitalization in Heart Failure With Reduced Ejection Fraction. Circ Heart Fail 2016;9.

35. Owan TE, Hodge DO, Herges RM, Jacobsen SJ, Roger VL, Redfield MM. Trends in Prevalence and Outcome of Heart Failure with Preserved Ejection Fraction. N Engl J Med 2006;355:251–259.

36. Rallidis LS, Makavos G, Nihoyannopoulos P. Right ventricular involvement in coronary artery disease: Role of echocardiography for diagnosis and prognosis. J Am Soc Echocardiogr 2014;27:223–229.

37. Haddad F, Doyle R, Murphy DJ, Hunt SA. Right ventricular function in cardiovascular disease, part II: Pathophysiology, clinical importance, and management of right ventricular failure. Circulation 2008;117:1717–1731.

38. Gennaro L De, Ferrara M, Bertini M. The spontaneous K-complex during stage 2 sleep: Is it the ‘forerunner’ of delta waves? Neurosci Lett 2000;291:41–43.

39. Halász P. K-complex, a reactive EEG graphoelement of NREM sleep: An old chap in a new garment. Sleep Med Rev 2005;9:391–412.

40. Bastien C, Campbell K. Effects of rate of tone-pip stimulation on the evoked K-Complex. J Sleep Res 1994;3:65–72.

41. Halász P. The K-complex as a special reactive sleep slow wave - A theoretical update. Sleep Med Rev 2016;29:34–40.

42. Johnson L, Karpan WE. Autonomic Correlates of the Spontaneous K-Complex. Psychophysiology 1968;4:444–452.

43. Gennaro L De, Gorgoni M, Reda F, Lauri G, Truglia I, Cordone S, Scarpelli S, Mangiaruga A, D’Atri A, Lacidogna G, Ferrara M, Marra C, Rossini PM. The Fall of Sleep K-Complex in Alzheimer Disease. Sci Rep 2017;7:39688.

44. Bruijn RFAG de, Ikram MA. Cardiovascular risk factors and future risk of Alzheimer’s disease. BMC Med 2014;12:1–9.

45. Himali JJ, Baril AA, Cavuoto MG, Yiallourou S, Wiedner CD, Himali D, Decarli C, Redline S, Beiser AS, Seshadri S, Pase MP. Association between Slow-Wave Sleep Loss and Incident Dementia. JAMA Neurol 2023;80:1326–1333.

46. Fultz NE, Bonmassar G, Setsompop K, Stickgold RA, Rosen BR, Polimeni JR, Lewis LD. Coupled electrophysiological, hemodynamic, and cerebrospinal fluid oscillations in human sleep. Science (80-) 2019;366:628–631.

47. Vijayakrishnan Nair V, Kish BR, Chong PL, Yang H-C (Shawn), Wu Y-C, Tong Y, Schwichtenberg AJ. Neurofluid coupling during sleep and wake states. Sleep Med 2023;110:44–53.

48. Bulcke L Van den, Peeters AM, Heremans E, Davidoff H, Borzee P, Vos M De, Emsell L, Stock J Van den, Roo M De, Tournoy J, Buyse B, Vandenbulcke M, Audenhove C Van, Testelmans D, Bossche M Van Den. Acoustic stimulation as a promising technique to enhance slow-wave sleep in Alzheimer’s disease: results of a pilot study. J Clin Sleep Med 2023;19:2107–2112.

49. Wunderlin M, Zeller CJ, Senti SR, Fehér KD, Suppiger D, Wyss P, Koenig T, Teunissen CE, Nissen C, Klöppel S, Züst MA. Acoustic stimulation during sleep predicts long-lasting increases in memory performance and beneficial amyloid response in older adults. 2023:1–11.

50. Oh HS-H, Rutledge J, Nachun D, Pálovics R, Abiose O, Moran-Losada P, Channappa D, Urey DY, Kim K, Sung YJ, Wang L, Timsina J, Western D, Liu M, Kohlfeld P, Budde J, Wilson EN, Guen Y, Maurer TM, Haney M, Yang AC, He Z, Greicius MD, Andreasson KI, Sathyan S, Weiss EF, Milman S, Barzilai N, Cruchaga C, Wagner AD, Mormino E, Lehallier B, Henderson VW, Longo FM, Montgomery SB, Wyss-Coray T. Organ aging signatures in the plasma proteome track health and disease. Nature 2023;624:164–172.

51. Helakari H, Korhonen V, Holst SC, Piispala J, Kallio M, Väyrynen T, Huotari N, Raitamaa L, Tuunanen J, Kananen J, Järvelä M, Tuovinen T, Raatikainen V, Borchardt V, Kinnunen H, Nedergaard M, Kiviniemi V. Human NREM Sleep Promotes Brain-Wide Vasomotor and Respiratory Pulsations. J Neurosci 2022;42:2503–2515.

52. Picchioni D, Özbay PS, Mandelkow H, Zwart JA de, Wang Y, Gelderen P van, Duyn JH. Autonomic arousals contribute to brain fluid pulsations during sleep. Neuroimage 2022;249:118888.

53. Bojarskaite L, Vallet A, Bjørnstad DM, Gullestad Binder KM, Cunen C, Heuser K, Kuchta M, Mardal KA, Enger R. Sleep cycle-dependent vascular dynamics in male mice and the predicted effects on perivascular cerebrospinal fluid flow and solute transport. Nat Commun 2023;14.

54. Baker FC, Driver HS. Circadian rhythms, sleep, and the menstrual cycle. Sleep Med 2007;8:613– 622.

55. Alzueta E, Zambotti M De, Javitz H, Dulai T, Albinni B, Simon KC, Sattari N, Zhang J, Mednick SC, Baker FC, Alzueta E, Zambotti M De, Javitz H, Dulai T, Albinni B, Simon KC, Sattari N, Zhang J, Shuster A, Sara C, Alzueta E, Zambotti M De, Javitz H, Dulai T, Albinni B. Symptoms Across the Menstrual Cycle with the Oura Ring in Healthy Women Tracking Sleep, Temperature, Heart Rate, and Daily Symptoms Across the Menstrual Cycle with the Oura Ring in Healthy Women. 2022.

56. Gómez-Santos C, Saura CB, Lucas JAR, Castell P, Madrid JA, Garaulet M. Menopause status is associated with circadian- and sleep-related alterations. Menopause 2016;23:682–690.

57. Renard Y, Lotte F, Gibert G, Congedo M, Maby E, Delannoy V, Bertrand O, Lécuyer A. OpenViBE: An open-source software platform to design, test, and use brain-computer interfaces in real and virtual environments. Presence Teleoperators Virtual Environ 2010;19:35–53.

58. Delorme A, Makeig S. EEGLAB: an open source toolbox for analysis of single-trial EEG dynamics including independent component analysis. J Neurosci Methods 2004;134:9–21.

59. Bigdely-Shamlo N, Mullen T, Kothe C, Su K-M, Robbins KA. The PREP pipeline: standardized preprocessing for large-scale EEG analysis. Front Neuroinform 2015;9:16.

60. Rogasch NC, Sullivan C, Thomson RH, Rose NS, Bailey NW, Fitzgerald PB, Farzan F, Hernandez-Pavon JC. Analysing concurrent transcranial magnetic stimulation and electroencephalographic data: A review and introduction to the open-source TESA software. Neuroimage 2017;147:934– 951.

61. Delorme A, Sejnowski T, Makeig S. Enhanced detection of artifacts in EEG data using higher-order statistics and independent component analysis. Neuroimage 2007;34:1443–1449.

62. Makeig S, Debener S, Onton J, Delorme A. Mining event-related brain dynamics. Trends Cogn Sci 2004;8:204–210.

63. Makeig S, Bell AJ, Jung T-P, Sejnowski TJ. Independent Component Analysis of Electroencephalographic Data. Proceedings of the 8th International Conference on Neural Information Processing Systems. Cambridge, MA, USA: MIT Press; 1995. p145–151.

64. Pion-Tonachini L, Kreutz-Delgado K, Makeig S. {ICLabel}: An automated electroencephalographic independent component classifier, dataset, and website. Neuroimage 2019;198:181–197.

65. Bernardi G, Siclari F. Local Patterns of Sleep and Wakefulness. Handbook of Sleep Research. Elsevier; 2019. p33–47.

66. Steriade M, Timofeev I, Grenier F. Natural Waking and Sleep States: A View From Inside Neocortical Neurons. J Neurophysiol 2001;85:1969–1985.

67. Behar JA, Rosenberg AA, Weiser-Bitoun I, Shemla O, Alexandrovich A, Konyukhov E, Yaniv Y. PhysioZoo: A novel open access platform for heart rate variability analysis of mammalian electrocardiographic data. Front Physiol 2018;9:1–14.

68. R Core Team. R: A language and environment for statistical computing. 2013.

69. Bates D, Mächler M, Bolker B, Walker S. Fitting Linear Mixed-Effects Models Using lme4. J Stat Softw 2015;67:1–48.

70. Vallat R. Pingouin: statistics in Python. J Open Source Softw 2018;3:1026.

