## Supplemental Material for "Cardiovascular responses to natural and auditory evoked slow waves predict post-sleep cardiac function"

Giulia Alessandrelli *et al.*

**Fig S1**

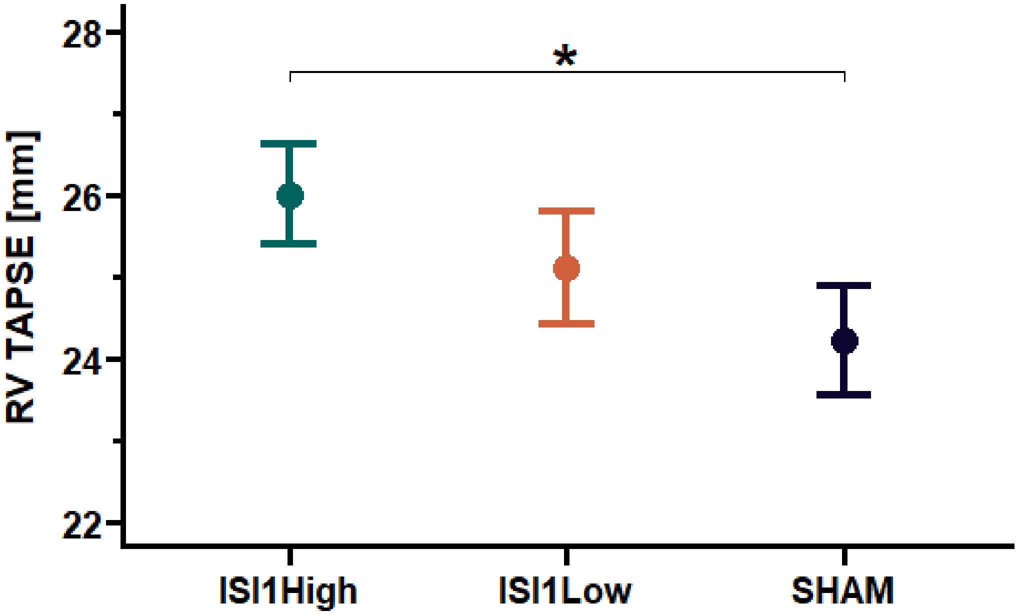


**Effects of auditory stimulation on echocardiography parameter estimating right-ventricular systolic function.** Right-ventricular tricuspid annular plane systolic excursion (TAPSE) is presented as mean ± standard error of the mean for n = 18 participants. P-values have been computed based on a linear-mixed effect model with the fixed factor Condition ($ISI1_{High}$, $ISI1_{Low}$, or SHAM) and random factor subject. Post-hoc comparison between conditions has been computed through paired t-tests and corrected for multiple comparisons by applying the Bonferroni correction. *: $p < 0.05$
